# Mapping of Glutamate Metabolism using 1H FID-MRSI after oral Administration of [1-^13^C]Glc at 9.4 T

**DOI:** 10.1101/2022.09.05.506579

**Authors:** Theresia Ziegs, Loreen Ruhm, Andrew Wright, Anke Henning

**Affiliations:** High-Field MR Center, Max Planck Institute for Biological Cybernetics, Max-Planck-Ring 11, 72076 Tübingen, Germany; IMPRS for Cognitive and Systems Neuroscience, Otfried-Müller-Str. 27, 72076 Tübingen, Germany; Advanced Imaging Research Center, University of Texas Southwestern Medical Center, 5323 Harry Hines Blvd., Dallas, Texas 75390, United States

**Keywords:** glutamatergic metabolism, [1-^13^C]Glc labeling, ultra-high field strengths, proton magnetic resonance spectroscopy imaging, human brain

## Abstract

Glutamate is the major excitatory transmitter in the brain and malfunction of the related metabolism is associated with various neurological diseases and disorders. The observation of labeling changes in the spectra after the administration of a ^13^C labelled tracer is a common tool to gain better insights into the function of the metabolic system. But so far, only a very few studies presenting the labelling effects in more than two voxels to show the spatial dependence of metabolism.

In the present work, the labeling effects were measured in a transversal plane in the human brain using ultra-short TE and TR 1H FID-MRSI. The measurement set-up was most simple: The [1-^13^C]Glc was administered orally instead of intravenous and the spectra were measured with a pure 1H technique without the need of a ^13^C channel (as Boumezbeur et al. demonstrated in 2004). Thus, metabolic maps and enrichment curves could be obtained for more metabolites and in more voxels than ever before in human brain. Labeling changes could be observed in [4-^13^C]glutamate, [3-^13^C]glutamate+glutamine, [2-^13^C]glutamate+glutamine, [4-^13^C]glutamine, [2,3-^13^C]aspartate and [6-^13^C]N-acetyl aspartate with a high temporal (3.6 min) and spatial resolution (32×32 grid with nominal voxel size of 0.33 μL) in five volunteers.

## Introduction

Abnormal glutamate (Glu) brain levels and function of the glutamatergic system have been found in different psychiatric disorders^1–4^ and neurological diseases including leukodystrophies and mitochondrial disorders^5^, epilepsy^6,7^and Huntington’s disease^8^ to just name a few. Carbon-13 Magnetic Resonance Spectroscopy (^13^C MRS) in combination with administering a ^13^C labelled substrate as tracer such as [1-^13^C]glucose (Glc) is a valuable tool to gain insight into brain metabolism. Different metabolic processes from glycolysis to the tricarboxylic acid (TCA) cycle and glutamatergic metabolism can thus be observed by following the incorporation of the ^13^C nucleus from Glc into different metabolites further downstream. The incorporation of the ^13^C nuclei into different metabolites were described in detail in previous publications^9–12^.

So far, mainly two different approaches were used to investigate the incorporation of ^13^C label into downstream metabolites: direct ^13^C and indirect ^1^H-[^13^C] MRS. Due to the enormous amount of related studies the reader is referred to review articles on these methods and applications^13–17^. Direct and indirect ^13^C methods both induce high specific absorption rates associated with heteronuclear decoupling and need special ^13^C hardware such as broadband amplifiers, multinuclear transmitter and receiver boards, dual-tune ^1^H/^13^C radiofrequency coils as well as non-standard sequences sending pulses on both ^1^H and ^13^C frequencies. Hence, Boumezbeur et al.^18^ proposed a new approach to measure metabolic rates. He showed the feasibility of conventional ^1^H MRS to measure metabolic rates without the use of any dedicated ^13^CMRS specific hardware. With this method, the incorporation of the ^13^C nuclei into different metabolites can be observed by the temporal changes in the spectral pattern in the ^1^H spectra due to heteronuclear scalar coupling of ^1^H and ^13^C, which for instance may lead to a splitting of singlet resonances into doublets^19^. To illustrate the effect: When ^13^C nuclei are incorporated into metabolites, the signal intensity of spectral peaks from the ^12^C-bonded protons decrease, while the peaks from ^13^C-bonded protons increase. The temporal changes of the spectra give insights into the speed of the corresponding metabolic processes.

So far, Boumezbeur’s method was applied in only a few studies in animal brain^20,21^ and recently also in human brain^7,22–24^. These studies measured the temporal changes in the MR spectra in a maximum of two voxels. In addition, most other studies using ^13^C or ^1^H-[^13^C] MRS techniques showed only data from one or two voxels. Only four studies measured the temporal changes of the ^13^C label incorporation in multiple volumes or slices using multivolume ^1^H-[^13^C] MRS^25^ or ^1^H-[^13^C] MRSI techniques in rat brain at 7T^26^ or ^1^H-[^13^C] MRSI techniques in human brain at 4T^27^ and 2T^28^. Nonetheless, the number of voxels from which data were measured were still rather small and/or the temporal resolution low: In the rat brain, de Graaf et al.^25^ measured 3 volumes within 6.4 min, Hyder et al.^26^ at least a slice with a grid of 16×16, but with a low temporal resolution of 10.7 min. In human brain, Watanabe et al.^28^ acquired data from a slice with 4 voxels with a nominal 37 ml volume with 15 min time resolution in the occipital lobe. While Pan et al.^27^ measured data from 16 voxels in a row in the coronal area with 4.5 min temporal resolution and a nominal voxel size of 6 ml. Both human MRSI studies acquired data from a small region in the occipital cortex. Pan et al. segmented the data and obtained rates of the tricarboxylic acid cycle in gray and white matter (GM, WM, respectively). Nonetheless, the investigated regions were rather small and hence no information about regional differences in the metabolism could be obtained. Especially, the frontal cortex would be of interest where MRS studies found more pronounced abnormalities in diseases than in the occipital lobe^29–31^.

The aim of this study was to show ^13^C label incorporation in different metabolites with regional and tissue type specificity using a substantially higher temporal and spatial resolution and spatial coverage than previous human studies. Moreover, metabolic maps as well as enrichment curves from a higher number of metabolites than the previous human spectroscopic imaging studies, which presented enrichment curves^27^ and maps^28^ for [4-^13^C]Glu only, was the aim. By applying ultra-short TE and TR single-slice 1H FID-MRSI at 9.4 T, data with high temporal (3.6 min) and spatial (0.7 ml) resolution could be obtained. Taking advantage of Boumezbeur’s technique in addition to the simple set-up of an oral [1-^13^C]Glc administration, metabolic maps as well as enrichment curves for [4-^13^C]Glu, [4-^13^C]glutamine (Gln), [3-^13^C]glutamate+glutamine(Glx), [2-^13^C]Glx, [2,3-^13^C]aspartate (Asp) and [6-^13^C]N-acetyl aspartate (NAA) could be determined. Due to the slice position in a transversal plane parallel to the corpus callosum, data from the frontal cortex could also be obtained, which was not done by the previous MRSI studies mentioned above.

## 2. Methods

The Glc preparation and the overall procedure of this experiment was very similar to a single-voxel spectroscopy (SVS) study published recently by the same group^24^. For clarity, the methods parts, which are the same as before were not rephrased.

Data and code can be made available upon request due to the huge size of MRSI data

### 2.1 Human Subjects

Labeling effects after the oral administration of [1-^13^C]Glc were measured in one slice above the corpus callosum in 5 healthy volunteers (2 female, 3 male, mean age 29±2 years). Before the measurement started, the volunteers gave their written informed consent according to the local research ethics regulations, the current version of the Declaration of Helsinki, DIN EN ISO 14 155 and were approved by the Institutional Review Board of the University of Tübingen.

### 2.2 [1-^13^C]Glc administration

The volunteers fastened for 9 hours before the measurement started. Before and after the scan the blood sugar level was tested with a glucometer (Accu-Check, Roche Diabetes Care GmbH, Mannheim, Germany) to detect possible hypoglycemia after the Glc administration. Hypoglycemia was not encountered for any subject. For each volunteer a solution containing 0.75g of [1-^13^C]Glc (Aldrich Chemical Company, Miamisburg, Ohio, USA; API for clinical studies) per kilogram body weight was prepared the subjects drank the Glc solution after the acquisition of the first MRSI slice. If possible, the volunteers drank the solution while lying down to keep the head as still as possible. The volunteers drank with a straw; since the straw was flexible, the volunteer could adjust the speed of solution entering the mouth easily.

### 2.3 Data Acquisition

All measurements were performed using a 9.4T Magnetom whole-body MR scanner (Siemens Healthineers, Erlangen, Germany) with an in-house built radiofrequency (RF) array coil with 18 transmit and 32 receive channels^32^. The subjects lay supine on the scanner table. Sagittal gradient-echo scout images were acquired for the positioning of the transversal slice (FOV 220 mm x 220 mm x 7 mm) just above the corpus callosum, see Figure 1a.

**Figure 1:**
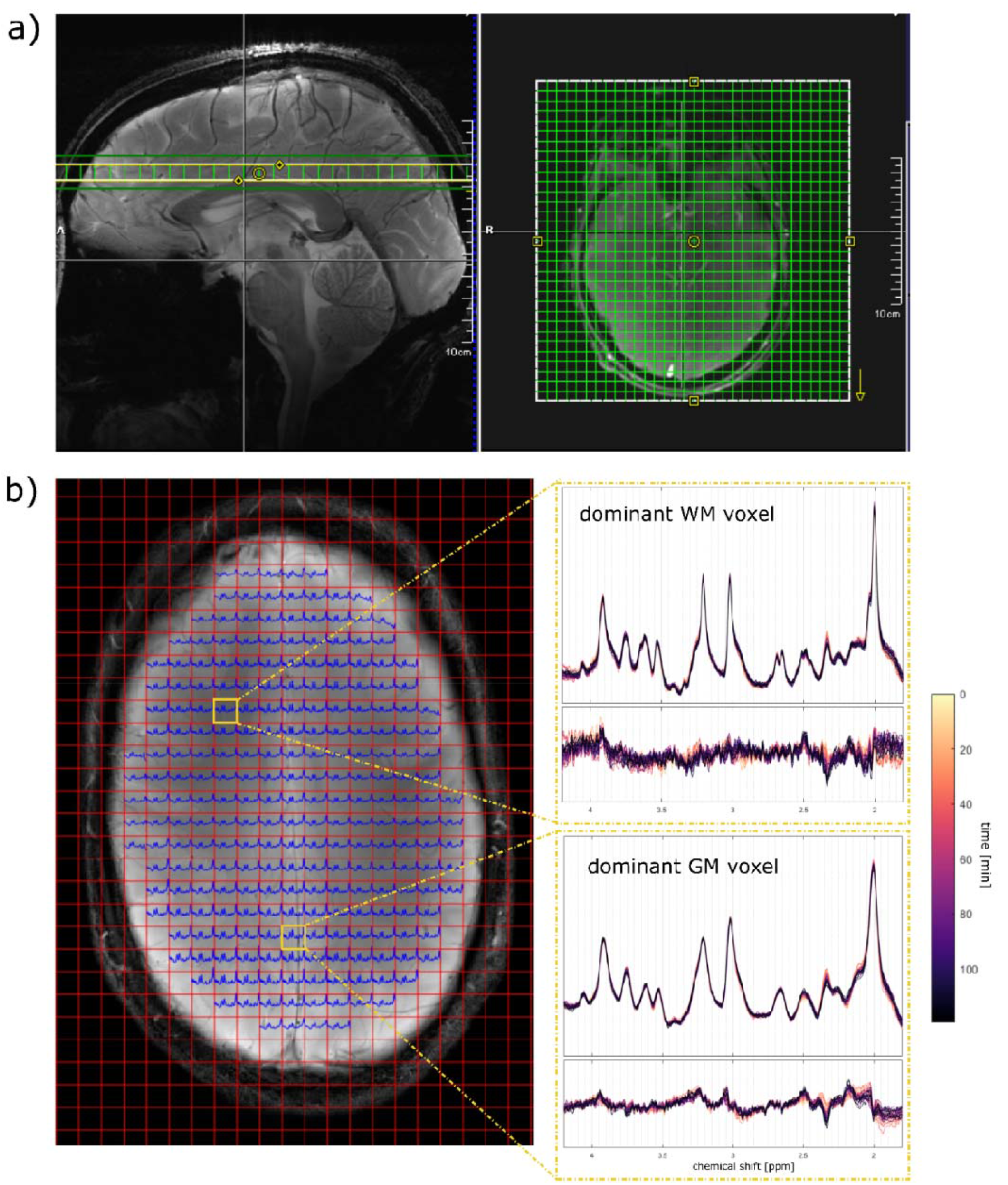
a) Position of the MRSI slice (220 mm x 220 mm x 7 mm; resolution 32×32) parallel to the corpus callosum (yellow box) and the corresponding shim volume (green box). b) Pre-Glc spectra for voxels within the brain (blue) and additionally, spectra and difference spectra for different time points from two voxel positions from the major gray and white matter area. Data from volunteer 1.

The slice was shimmed using the vendor implemented image-based second-order B_0_ shimming routine with a shim volume of 220 mm x 220 mm x 15 mm with the same center as the MRSI slice. A customized ^1^H FID MRSI sequence^33,34^ (resolution 32×32 with elliptical k-space shuttering, TR = 300 ms, TE*=1.5 ms, flip angle = 48°, spectral width= 8000 Hz, acquisition time = 128 ms) with optimized water suppression^34^ and without outer volume or lipid suppression was acquired. Subsequently, a non-water suppressed reference scan with the same resolutions and scan parameters as the ^1^H MRSI scan was measured. The scan time for one ^1^H MRSI acquisition was 3.6 minutes.

After these initial measurements were executed, the scanner table was pulled out and the volunteers drunk the Glc solution as fast as possible. After this short break, the scanner table was pushed back to its previous position inside the bore and a localizer was applied to verify the positioning of the slice. In case of a changed head position, the gradient-echo scout sequences to relocate the MRSI volume at the exact same position as prior to the Glc intake and B_0_ shimming was repeated before continuing to apply the ^1^H MRSI sequence. If no repositioning of the voxel was needed (4 of the 5 measurements), ^1^H MRSI data acquisition was continued immediately. The time after the Glc administration and the start of the ^1^H MRSI measurement was 4.5-12 minutes. As many as possible of the water suppressed ^1^H MRSI acquisitions were repeated to fill a maximum total scan time of 2 hours (21-35 spectra were acquired within 70-120 minutes).

In a different session, an MP2RAGE from each volunteer was acquired for gray and white matter segmentation.

### 2.4 Data Processing

^1^H MRSI raw data was processed with in-house-written code in Matlab (version 2016a; MathWorks, Natick, MA) as also described before in Nassipour et al^34^ including spatial reconstruction by fast fourier transformation, eddy current correction^35^, coil combination using singular value decomposition (SVD), water removal with Hankel SVD, missing point prediction and phase correction. The MP2RAGE images were reconstructed^36^ and segmented using the SPM12 Matlab package.

### 2.5 Data Fitting

The reconstructed MRSI data was fitted using LCModel (V6.3-1L, the control file can be found in the Supplementary Material)^37^ with a basis set simulated with VeSPA (version 0.9.5 https://scion.duhs.duke.edu/vespa/^38^) consisting of the following basis spectra: aspartate (Asp), γ-aminobutyric acid (GABA), glycine, glutathione (GSH), myo-Inositol (mI), NAAG, scyllo-inositol (scyllo), taurine (Tau), Glc, Glu, Gln, NAA, total creatine (tCr), total choline (tCho) and a simulated macromolecular baseline (MM). Since changes in the spectral pattern due to ^13^C label incorporation are detectable via decreasing peaks of the ^12^C-bonded proton signals the Glu, Gln, and NAA spectra were modified: the Glu and Gln spectra were splitted into its individual resonances to account for labeling in different ^13^C positions: [4-^12^C]Glu, [4-^12^C]Gln, [3-^12^C]Glx, [2-^12^C]Glx. These peaks are subsequently designated as Glu4, Gln4, Glx3 and Glx2, respectively, according to the number of the ^13^C position. Furthermore, NAA was splitted into its acetyl (^2^CH_3_) and aspartate (^2^CH) moiety to account for peak changes due to the labeling of the acetyl group, which is further denoted as NAA6 (6^th^ carbon position). Since also Asp becomes labeled at the 2^nd^ and the 3^rd^ carbon position, it will be called Asp2,3. Splitting of these Asp peaks did not yield consistent fitting results. Furthermore, tCr was splitted too into its CH_3_ moiety peak at approx. 3 ppm (tCr CH_3_) and the CH_2_ moiety. The temporal changes in the CH_3_ peak serves as a proof of spectral stability. The fit of all metabolites from one voxel is shown in Supplementary Figure S1 for the first time point and the fit of those metabolites, which change over time, are shown in Supplementary Figure S2. The peaks of the spectra from single voxels of the MRSI slice were too broad to account for increasing satellite peaks from the ^13^C-bonded protons as it is detectable in summed difference spectra, see Figure 2c, and as it was done in 1H SVS studies^18,23,24^.

**Figure 2:**
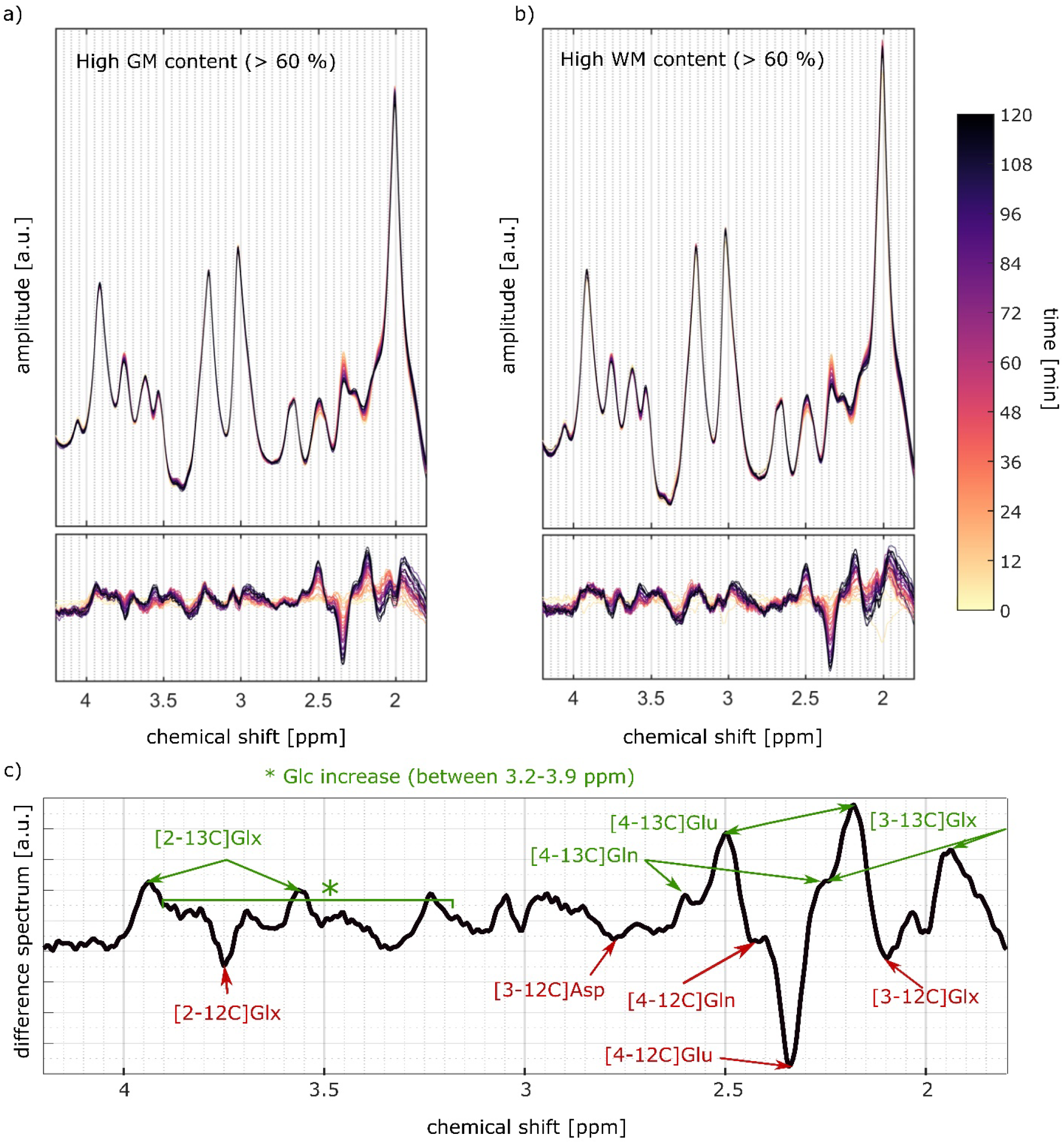
Temporal series of summed spectra for voxels with a) high gray matter and b) white matter content (>60 %) with the corresponding difference spectra. Colors indicate the measured time points. C) Averaged last difference spectrum from high gray matter voxels with decreased metabolite peaks indicated in red, and increased metabolite peaks indicated in green as far as determination of the corresponding metabolites were possible. Data from volunteer V1.

J-coupling constants and chemical shifts were taken from Govindaraju et al.^39,40^ except for the J-coupling constants for GAB, which were taken from Near et al^41^. The line broadening of the basis spectra was set to 3 Hz. The spectra were fitted between 1.8 and 4.2 ppm. The simulated MM baseline (MM_AXIOM_ in Wright et al.^42^) was used since the acquisition of a matching experimental macromolecular spectra was not possible with the same short TR due to SAR constraints of the required double inversion recovery sequence. The simulated MM spectrum used in this work considered T_1_- and T_2_-relaxation times of MM molecules as well as the linewidths of the MM peaks^43,44^ and was successfully used in ^1^H MRSI data before^45,46^.

### 3.6 Percent Enrichment Calculation

The median of the peak intensities for ^13^C labeled metabolites from all voxels, from voxels with high GM (> 60 %) and voxels with high WM (> 60 %) content was calculated for each volunteer separately. The temporal changes of the median curves were fitted with an exponential function *y*(*time*) =*a* + *b* · *e*^-|*k*|·*time*^. The corresponding ^13^C percent enrichments (PEs) were calculated with: 1 − *y*(*time*)/*y*(*time* = 0). The PE curves were averaged across volunteers. Finally, a two-sided Wilcoxon rank sum test was performed to test for significant differences in the mean final PE values from voxels with high GM content to those with high WM content at the 5% significance level.

## 3. Results and Discussion

### 3.1 Labeling Effects on Spectral Pattern

The averaged GM and WM spectra and respective difference spectra for all time points show systematic changes in the chemical shift region between 2 ppm – 4 ppm, see Figure 2a+b. Figure 2c illustrates that these spectral pattern changes arise from ^13^C labeling of Glu, Gln, Asp and the increase of the sum of ^12^C and ^13^C labelled Glc concentration. These changes are consistent to results shown in in 1H SVS data at 9.4T after oral Glc administration^24^.

### 3.2 Metabolic Maps and Enrichment Maps

Concentration ratio (/tCr) maps for ^12^C labelled Glu4, Glx3 and Glx2 for different time steps shows that the concentration decrease of the ^12^C-bonded protons decrease visually across the entire brain slice, see Figure 3b. tCr ratios were used to include the correction for the transmit (B1^+^) and receive (B1^-^) radiofrequency field correction^34^. The PE map of Glu4 (2h after the Glc intake) reveal a dependence of label increase on tissue types, see Figure 3a+c.

**Figure 3:**
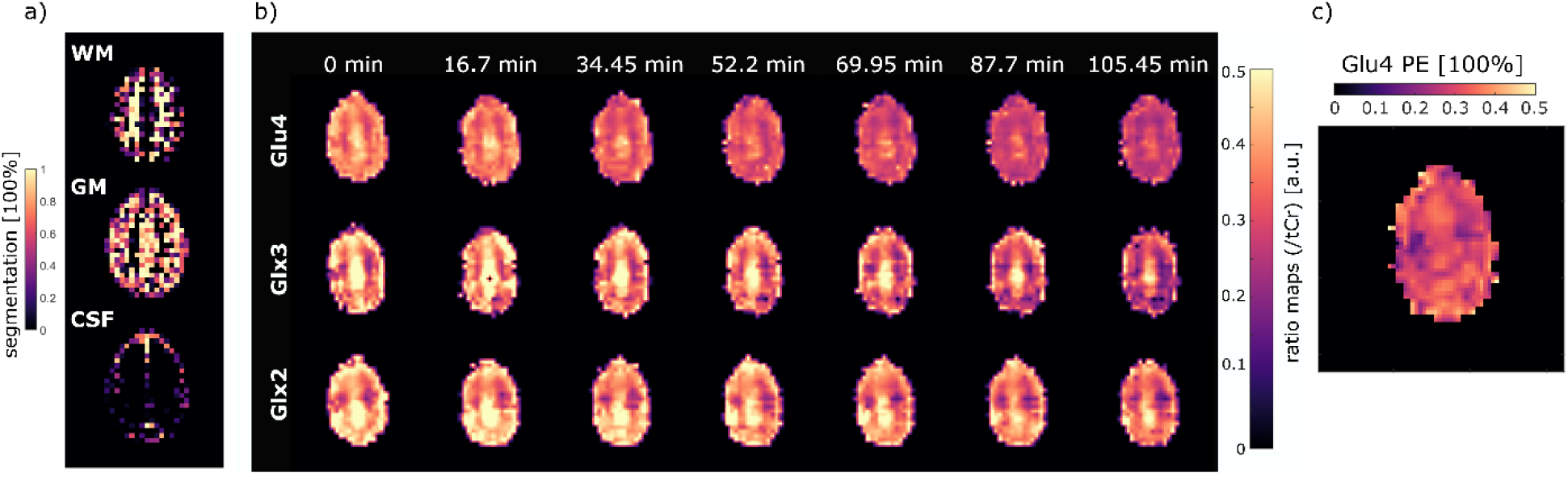
White matter, gray matter and CSF segmentation of the MRSI slice. Ratio maps (/tCr) for different time points of Glu4, Glx3, Glx2 and the corresponding relative changes to the first MRSI data set in %; data from volunteer 1.

### 3.3 Time Courses of Labeled Metabolites

The tissue dependence of the label effects in different metabolites are seen in the mean PEs averaged across all volunteers for voxels with high GM (>60%) or WM (>60) content, see Figure 4 along with the 0.95 standard deviation interval.

**Figure 4:**
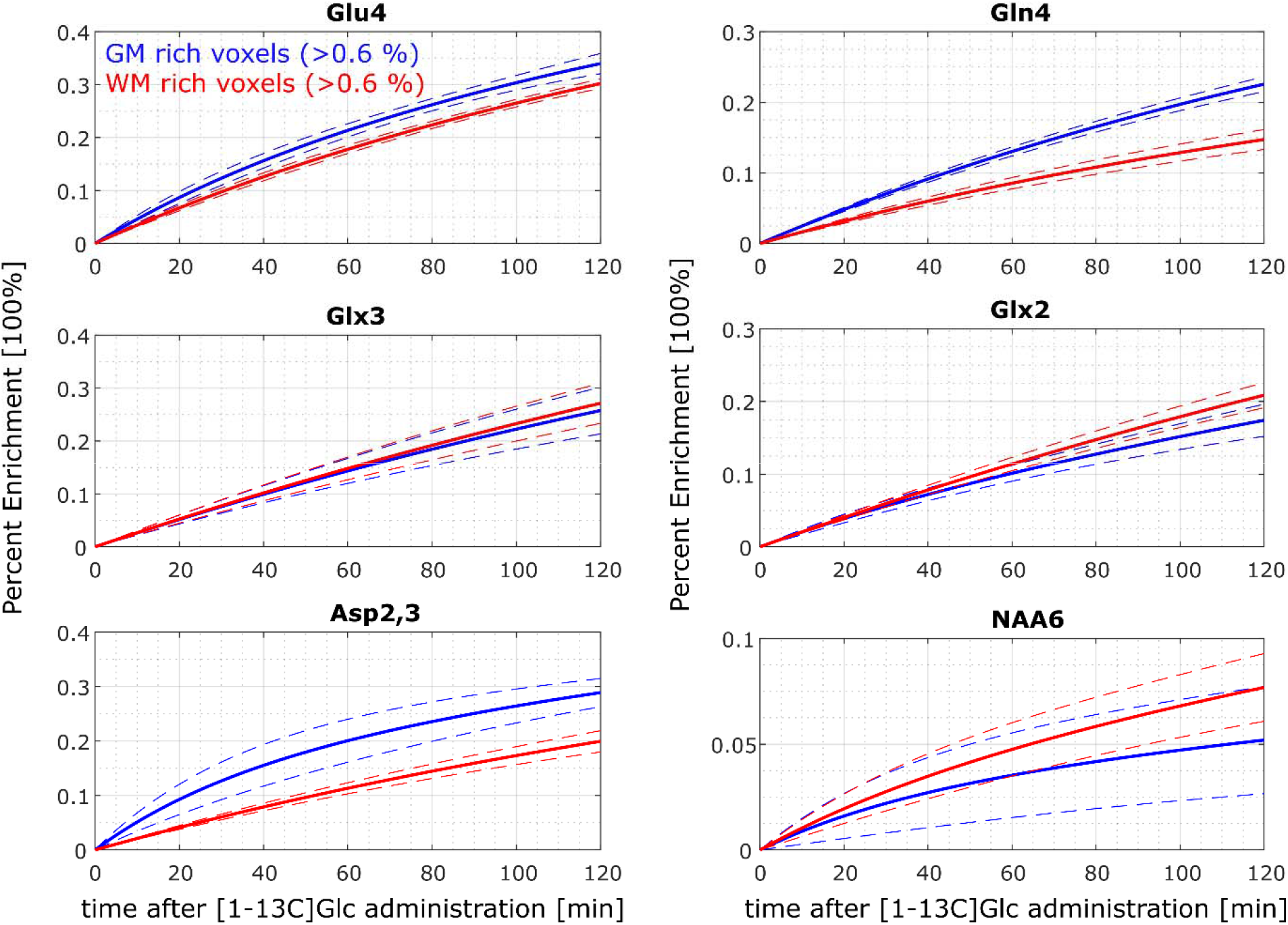
Mean percent enrichment for different metabolite peaks from voxels with high gray matter or white matter content (>60%, blue and red, respectively). Mean (solid line) and 0.95*std intervall (dashed line) across all volunteers.

The PE after 2h for all voxels and those with high GM and WM (>0.6%) are collected in Table 1: Two hours after the Glc administration the Glu4 peak was reduced by 32±2 % (mean ± standard deviation across subjects); the Gln4 peak by 17±5 %; Glx3 by 27±9 %; Glx2 by 18±4%; Asp2,3 by 26±5%; NAA6 by 8±4. The differences in WM and GM voxels are found to be significant in Glu4, Gln4 and Asp2,3.

**Table 1:** Percent enrichment after 2h of measurements averaged across subjects (mean, standard deviation) for voxels with high GM and WM content (>60 %) and all voxels within the brain slice. Wilcoxon rank sum test was performed to test the significance for differences in the mean final PE values for dominant GM and WM voxels 2h after the Glc intake.

| PE<br>[%] | GM |  | WM |  | ALL |  | Significance |
| --- | --- | --- | --- | --- | --- | --- | --- |
|  | mean | std | mean | std | mean | std |  |
| **Glu4** | 34 | 4 | 30 | 2 | 32 | 2 | 1 |
| **Gln4** | 23 | 2 | 15 | 3 | 17 | 5 | 1 |
| **Glx3** | 26 | 10 | 27 | 8 | 27 | 9 | 0 |
| **Glx2** | 17 | 5 | 21 | 4 | 18 | 4 | 0 |
| **Asp2,3** | 29 | 5 | 20 | 4 | 26 | 5 | 1 |
| **NAA6** | 5 | 5 | 8 | 3 | 8 | 4 | 0 |
| **Cr CH<sub>3</sub>** |  |  |  |  | 0.2 | 3 |  |

The single volunteer’s time courses for the same metabolites and tCr CH_3_ can be found in the Supplementary Figure S3 and demonstrate reproducibly high data quality and consistency. The tCr CH_3_ peak by maximum 2 % after 2h of measurement, except for volunteer V4, which increases by approx. 3 %. Although, the time course from this volunteer shows unexpected behavior in the first hour, the change of tCr is within the error margin and the mean PEs across all volunteers would not change significantly when excluding volunteer V4.

Comparing the present results to the literature is difficult, since spectroscopic imaging has only been rarely used to overserve the spatial dependence of metabolism after the administration of a ^13^C labeled substrate either in animals^25,26,47,48^ or humans^27,28^. In human brain, Pan et al.^27^ showed PE curves for Glu4 with a saturated PE of approx. 40% in coronal slice with 16 volumes (6 mL each) after the infusion of [U-^13^C]Glc using indirect short spin echo single quantum ^13^C editing with a time resolution of 4.5 min. Watanabe et al.^28^ presented Glu4 peak height curves as well as temporal changes in the Glu4 maps but no PEs for 4 volumes (37 mL each) in a slice in the occipital lobe using also an indirect method (heteronuclear single quantum coherence, HSQC) by administering [1-^13^C]Glc orally with different doses with 15 min time resolution.

The PE of Glu4 from the present study is slightly higher than the values after 2 to 3h (when saturation is expected) from other oral studies administering the same amount of [1-^13^C]Glc (20-24%^24,49^) or less [1-^13^C]Glc (16% using 0.65 g Glc/kg body weight instead of 0.75 g/kg^50^). Most studies use i.v. administration and showed maximum Glu4 PEs of 11-40%^23,27,49–55^, with our results being in the upper range of these values (literature is collected in Supplementary Table S1). Results from studies using oral intake should be cautiously compared to studies using i.v. administration due to the following limitations: First, the maximum PE is obviously depending on the amount of Glc given. While the oral intake contains a defined amount of Glc, the i.v. Glc administration procedure includes often a maintenance dose, whose amount is in many cases not reported. Secondly, the same amount in an oral as well as an i.v. administration, does not lead to similar maximum enrichments^50^. Third, the oral administration leads to delayed enrichments curves due to the gastrointestinal Glc absorption^5,49^. Thus, the saturation plateau would only be reached after 3h of measurement in contrast to the i.v. protocol, where the maximum PE is reached after 2h of measurement^49^. Unfortunately, 3h of measurement could not be performed in the present study due to ethical regulations related to the maximum scan time. Additionally, such a long scan time would be uncomfortable for the volunteers and increase the risk of motion. An additional break taking the subject out of the scanner – as suggested from Mason et al.^49^ – would entail slice-repositioning problems and was thus not realized in this study. Due to the different Glc dose and the additional route through the body, the PE values from i.v. studies cannot be directly compared to the oral results, but are still included into this manuscript since it is the far more common procedure.

Previously, it has been shown that Glu4 is labeled before Gln4^51,56^. In addition, the PE of Glu4 and Gln4 is found to be faster in GM than WM^25–27,55^ due to faster metabolism of the corresponding chemical reactions in GM than in WM^25–27,55^. Both is also found in the present data and the changes are significant at the 5 % significance level. Unfortunately, most publications did not compare GM and WM regions or did not segment their voxels at all. Lebon et al.^57^ found an enrichment ratio of Gln4/Glu4 to be 0.63 in a voxel containing 57 % GM using ^13^C MRS, which is similar to the ratio of 0.68 for the GM rich voxels (>60 %) in the present study. The enrichment of Gln4 is in the range of previous oral (approx. 15% in the occipital lobe^24,49^) and i.v. studies (10-25%^49,52,53^). The herein presented Glx3 values are at the upper range of previous values for oral administration (17-19 % after 2h^24^) or i.v. protocols (5-10%^23^ and 23%^53,54^). There is no significant change of the PEs in GM or WM voxels. The Glx2 is in the range of previous values (10-15 % after 1h^52^ and 13-23% after 2h^24^). In the present data, Glx2 is labeled slightly faster in WM voxels than in GM voxels, but the changes are not significant.

Asp enrichment is rarely reported in literature. In this study, Asp2 and Asp3 were not fitted separately since this led to ambiguous results of the Asp3 PEs. Nonetheless, the enrichment should be relatively similar because both are labeled in the first TCA turn^12^. The labeling through PC (3-10 % of the Glc consumption^58^) could result in slightly different maximum enrichments of Asp2 and Asp3, which could not be determined with the present approach. Previous studies using oral administration reported 50-100 % higher values for Asp3 enrichment than the present data^24^, but those values should be treated with caution due to artifacts mentioned in^24^. I.v. studies observing Asp3 enrichment found values similar to the Glu4 enrichment^51^ and values of approx. 27 %^54^, which is similar to herein reported Asp2,3 enrichment in the GM rich voxels. 12% PE was found for Asp2^10^, which is definitely lower than the herein presented values for Asp2,3. The faster increase of Asp2,3 in GM than WM can be again explained by the faster TCA cycle rate in GM^25–27,55^.

Although NAA6 enrichment is slow^12,51^, it was detectable in mouse and rat brain^12,59–61^. Respective data^12^ indicates an enrichments of approx. 5-10 % after 2h, which is similar to the values in the present study. No other human study showed NAA6 labeling, but labeling in NAA2 could not be found after approx. 1h^51^ or changes of 2 % after 2h^10,51^. NAA2 should be labeled subsequent to NAA6 since it is labeled later in the respective pathways^12^. The variance in the enrichments in WM and GM are too high to distinguish differences for both tissue types.

### 3.4 Limitations

In the present study, it was not possible to calculate metabolic rates since the brain or blood ^13^C Glc enrichment was not accessible. While total Glc levels as well as unlabeled and labelled Glc fractions are in principle measureable in ^1^H SVS data^24,62^, the line width in ^1^H MRSI data is too broad to measure (total) Glc levels in the spectra. A solution would be to sample the blood during the measurement as it was done in most other studies^49^, which was not possible in the current study inside a non-clinical setting.

The sensitivity could be improved by the use of [1,6-^13^C]Glc or [U-^13^C]Glc instead of Glc labeled at a single position, which doubles the sensitivity since both compounds lead to labelling of two pyruvate molecules instead of only one. However, it would be (2-5 fold) more expensive to perform such a study in humans, which is not affordable since the costs were already approx. 5000 euro/person.

Another problem of the present study set-up was the short break for drinking the Glc solution after the baseline ^1^H MRSI measurement. This break causes uncertainties in the relocation of the ^1^H MRSI slice. Although the position of the post-Glc ^1^H MRSI slice was compared visually to the pre-Glc MRSI slice, the possibility of a mismatch between both slice positions remain. The error is of minor importance since the enrichment curves were calculated using an exponential fit using all time points; so, possible errors in the first value are negligible. Only the mismatch of the B_0_ shim volume could cause imperfect shim results, but since the shim volume was chosen to be larger than the slice, it is probably of minor importance, too. A possible solution would be automatized repositioning of the 1H MRSI slice, which was not available on the present scanner.

In further studies, it would be interesting to use this technique for the observation of different labeling effects in different brain regions, e.g. frontal cortex vs. occipital lobe.

## 5. Conclusion

The present study demonstrates for the first time that glutamatergic metabolism can be derived from 1H MRSI data with high temporal and spatial resolution without the need of specialized ^13^C hardware or scan software at 9.4 T. Enrichment curves of Glu4, Gln4, Glx3, Glx2, Asp2,3, and NAA6 and enrichment maps of Glu4 have been obtained and revealed the possibility of this approach to investigate the dependence of the labeling effects on regions and tissue types across the brain.

## Abbreviations

FID: free induction decay
TE: echo time
TR: repetition time
MRSI: Magnetic Resonance Spectroscopic Imaging
Glc: glucose
Glu: glutamate
Gln: glutamine
Asp: Aspartate
PC: pyruvate carboxylase
TCA: tricarboxylic acid cycle
PDH: pyruvate dehydrogenase complex
SVS: single voxel spectroscopy

