## Supplementary for "Mapping of Glutamate Metabolism using 1H FID-MRSI after oral Administration of [1-^13^C]Glc at 9.4 T"

Supplementary material:

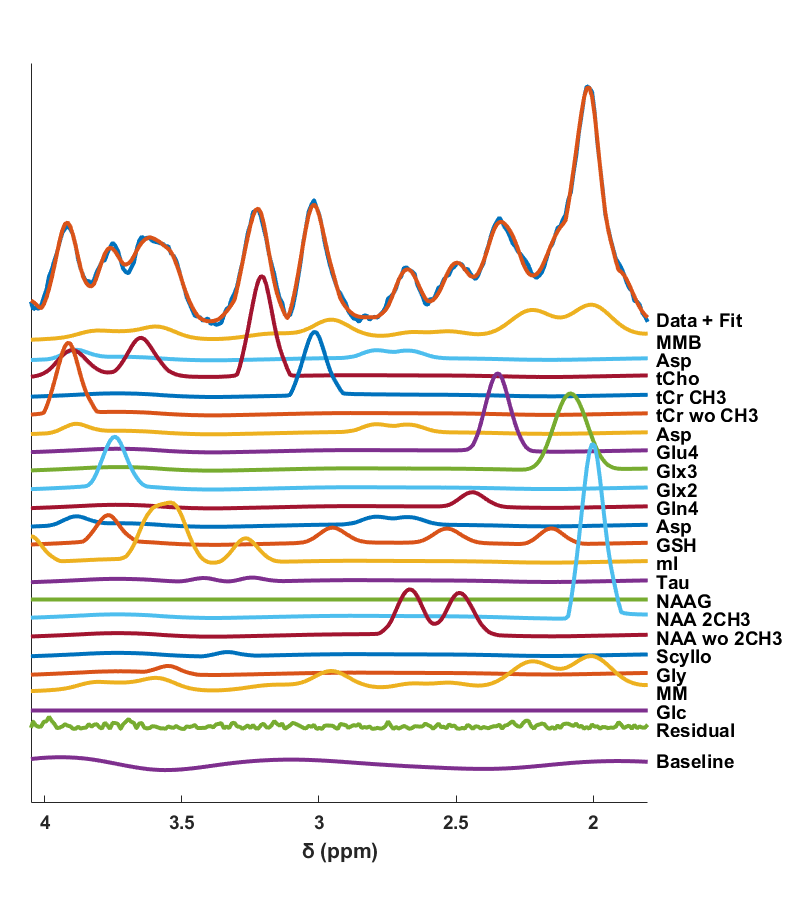

Figure S1: Data, residual and fit of all metabolites for a sample voxel from V1 in the middle of the brain for the first measurement.

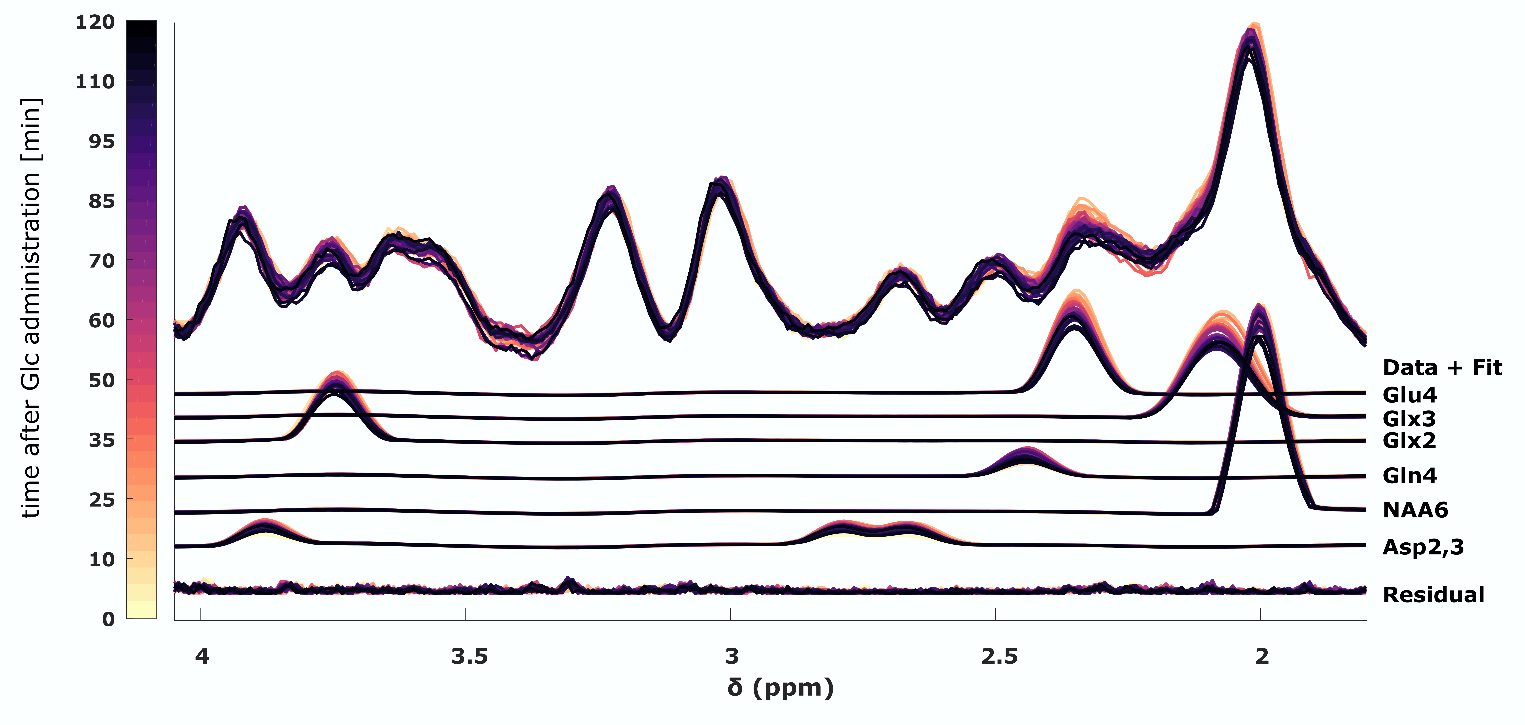

Figure S2: Data, fit and residual for a time series of spectra from volunteer V1 in the middle of the brain at position 16x16. Colors indicate different time points after Glc administration.

LCModel *.control file

$LCMODL

hzpppm= 399.719

deltat= 1.2500e-04

nunfil= 1036

neach= 50

nratio = 0

ppmend= 1.8

ppmst= 4.1

doecc= F

sddegp= 0

sddegz= 0

dows= T

wconc = 40873

atth2o = 1

dkntmn= 0.15

nsimul = 0

ncombi = 18

ndslic= 1

ndrows= 32

ndcols= 32

islice= 1

irowst= xi

irowen= xi

icolst= yi

icolen= yi

chcomb(17) = 'Glu2+Glu3+Glu4'

chcomb(18) = 'Gln2+Gln3+Gln4'

$END

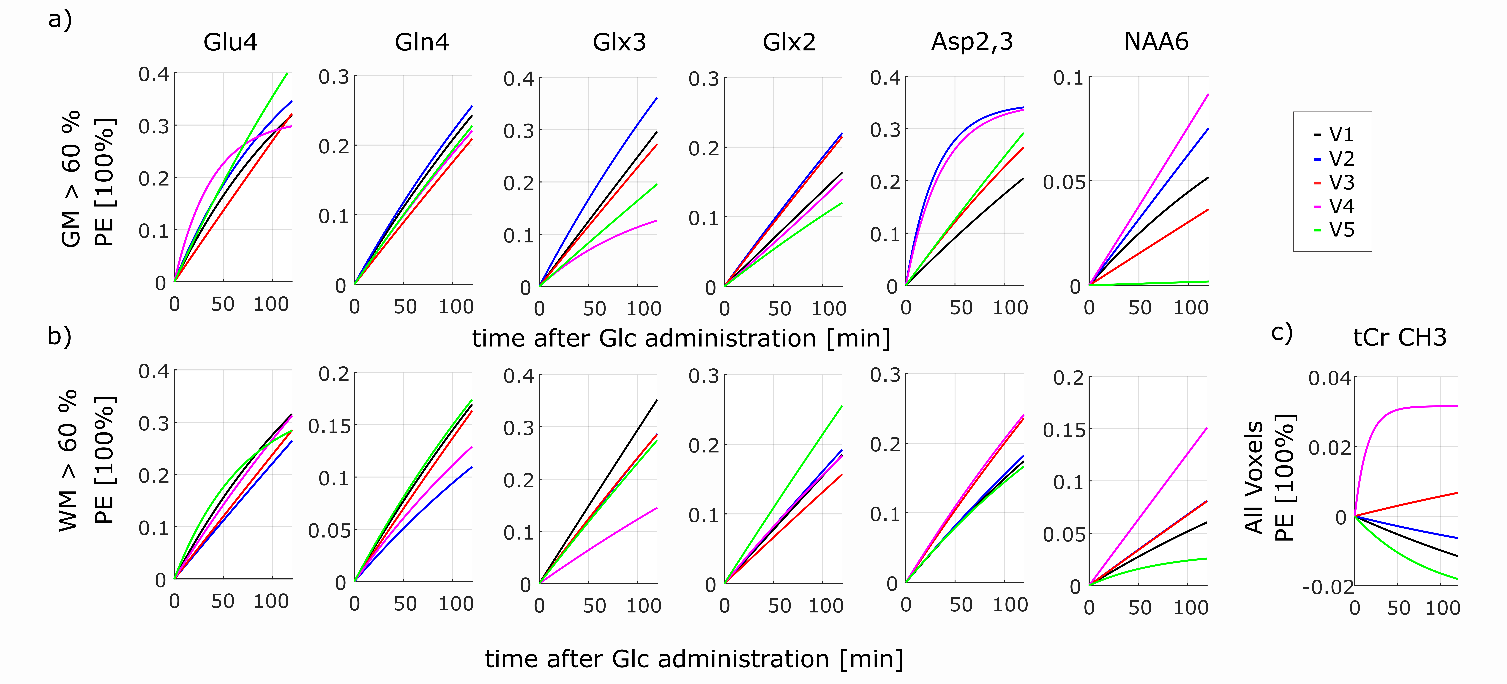

Figure S3: Inverse exponential fit of the median time courses of Glu4, Gln4, Glx3, Glx2, Asp2,3, NAA6 averaged over all voxels with a) GM > 60 % and b) WM > 60 %. Inverse exponential fit of the median time course of Cr CH3 averaged over all voxels. Volunteers V1-V5 are indicated with colors.

|  | oral/  i.v. | Labeled  Glc | B0 [T] | Method | Region | Segmentation | Glu4  [%] | Gln4  [%] | Glx3  [%] | Glx2  [%] | Asp2,3  [%] | NAA6  [%] |
| --- | --- | --- | --- | --- | --- | --- | --- | --- | --- | --- | --- | --- |
| present study | oral | 1-13C | 9.4 | 1H MRSI | transversal slice | WM rich voxels (>60 %) | 30* | 15* | 27* | 21* | 20* | 8* |
|  |  |  |  |  |  | GM voxels (>60 %) | 34* | 23* | 26* | 17* | 29* | 5* |
| Pan et al.  2000 | i.v. | U-13C | 4 | 1H-[13C]MRSI | coronal slice in the occipital lobe | yes | 40 |  |  |  |  |  |
| Ziegs et al. 2022 | oral | 1-13C | 9.4 | 1H MRS | occipital lobe | no | 24* | 15* | 19* | 19* | 41* (Asp3) |  |
|  |  |  |  |  | frontal cortex |  | 24* | (20*) | 17* | 23* | 37* (Asp3) |  |
| Mason et al. 2003 | oral | 1-13C | 2.1 | 13C MRS | occipital lobe? | no | 20 | 17 |  |  |  |  |
|  | i.v. |  |  |  |  |  | 18 | 10 |  |  |  |  |
| Moreno et al. 2001 | oral | 1-13C | 1.5 | 13C MRS | occipitoparietal region | mixed voxel | 16* |  |  |  |  |  |
|  | i.v. |  |  |  |  |  | 11-13 |  |  |  |  |  |
| Dehghani et al. 2020 | i.v. | 1-13C | 3 | 1H MRS | anterior cingulate cortex, posterior cingulate cortex | no | 15† |  | 5-10† |  |  |  |
| Bartnik et al. 2017 | i.v. | U-13C | 3 | 1H MRS | mesis temporal lobe | 37 % GM, 60 % WM | -27† Glu decrease |  |  |  |  |  |
| Moreno et al. 2001 | i.v. | 1-13C | 1.5 | 13C MRS | occipital region | No |  |  |  |  | 12 (Asp2) |  |
| Cchina et al. 2001 | i.v. | 1-13C | 3 | 13C MRS | occipital lobe | No | 30† | 25† |  | 15† (Glu2, Gln2) |  |  |
| Shen et al. 1999 | i.v. | 1-13C | 2.1 | 13C | occipital and  parietal lobe | no | 25 | 25 |  |  |  |  |
| Mason et al. 1999 | i.v. | 1-13C | 4.1 | 1H-[13C] MRS | cingulate sulcus | GM voxel | 19 |  |  |  |  |  |
|  |  |  |  |  | centrum semiovale | WM voxel | 23 |  |  |  |  |  |
| Gruetter et al. 1998 | i.v. | 1-13C | 4 | 13C MRS | occipital lobe | no |  |  |  |  | comparable Glu4 (Asp3) |  |
| Mason et al. 1995 | i.v. | 1-13C | 2.1 | 13C MRS | occipitoparietal region | GM voxel | 27 | 25 | 25 (Glu3) |  |  |  |
| Gruetter et al. 1994 | i.v. | 1-13C | 2.1 | 13C MRS | occipitoparietal region | No | 27 | 26 | 23 (Glu3) |  | 27 (Asp3) |  |

Table S1: Literature values of the label increase in different metabolites administering labeled Glucose. Only data from human studies listed. If saturation PE was not available, the maximum enrichment after 2h (*) or 1h (†) are shown, especially for most oral studies, the saturation plateau was not reached within the measurement time. If the publication did not directly mention the PE values, there were mostly taken from the presented curves. For some entries, the PE values were calculated using unlabeled and labeled metabolite concentration mentioned in the studies. Many studies show uptake values in arbitrary units or mM or report metabolic rates instead. Those were not listed here. After the own results, literature using MRSI and then studies using oral administration are listed. Afterwards, the paper articles are ordered backward in time.
